# Deep neural networks identify context-specific determinants of transcription factor binding affinity

**DOI:** 10.1101/2020.02.26.965343

**Authors:** An Zheng, Michael Lamkin, Cynthia Wu, Hao Su, Melissa Gymrek

## Abstract

Transcription factors (TFs) bind DNA by recognizing highly specific DNA sequence motifs, typically of length 6-12bp. A TF motif can occur tens of thousands of times in the human genome, but only a small fraction of those sites are actually bound. Despite the availability of genome-wide TF binding maps for hundreds of TFs, predicting whether a given motif occurrence is bound and identifying the influential context features remain challenging. Here we present a machine learning framework leveraging existing convolutional neural network architectures and state of the art model interpretation techniques to identify, visualize, and interpret context features most important for determining binding activity for a particular TF. We apply our framework to predict binding at motifs for 38 TFs in a lymphoblastoid cell line and achieve superior classification performance compared to existing frameworks. We compute importance scores for context regions at single base pair resolution and uncover known and novel determinants of TF binding. Finally, we demonstrate that important context bases are under increased purifying selection compared to nearby bases and are enriched in disease-associated variants identified by genome-wide association studies.

## Introduction

Binding of transcription factors (TFs) is one of the major transcriptional regulation mechanisms. TFs recognize and bind specific DNA sequences to control transcription, forming a complex system that guides gene expression^1^. Disruption of TF binding is the root cause of a wide range of prevalent diseases^2^, including cancer^3,4^, autoimmune disorders^5^, and cardiovascular diseases^6^. Thus, understanding the mechanisms controlling TF binding can provide insights into disease processes, eventually leading to potential treatments.

TFs typically recognize short specific motifs of 6-12bp^1^. However, these motifs cannot completely explain TF binding affinity. There is often only a partial overlap between experimentally determined binding sites in the genome and sequences matching the motif for a particular TF^1^. For example, in our study, we found that the SP1 TF motif occurs more than 3.6 million times across the human genome, whereas less than 1% of these occurrences are actually bound in a human lymphoblastoid cell line. Whether a particular motif instance is bound depends on multiple facets, including the accessibility of chromatin^7,8^, nucleosome positioning^9,10^, cooperative and competitive binding with other factors^11^, local GC content^10,12^, and local DNA tertiary structures^13^. Furthermore, these features and their importance for binding may vary greatly across cell types, loci, or development timepoints^1^. Studies have shown that many of these features are related to sequence context in the immediate vicinity of the TF motif itself^14–16^, implying that TF binding may be predicted directly from sequence information.

Several machine learning methods^17–24^ have proven successful in predicting TF binding from sequence. Many of these methods, such as DeepSEA^18^ and DanQ^20^, rely on convolutional neural networks (CNNs), which infer important sequence features in the binding regions and learn connections between TF binding and combinations as well as orientations of these sequence features. However, these CNN-based frameworks face several limitations. First, they use a multi-class training procedure focusing on open chromatin regions that are active in at least one cell type of interest, but do not contain regions inactive in all cell types as controls. Thus, their models cannot learn general features that distinguish bound vs. unbound genomic regions. Second, while these models have shown excellent prediction accuracy for a variety of marks and cell types, interpreting CNNs to derive meaningful biological insights remains challenging.

Here, we expand on existing CNN models, DeepSEA and DanQ, to develop a framework for predicting whether a particular instance of a TF will be bound and provide nucleotide-resolution insight into the context sequences essential for TF binding. By conditioning on sequences that contain the core TF motif in both the case and control samples, we allow our framework to discover sequence context features that determine binding status. In our framework, we use a transfer learning scheme to reduce the amount of data needed for training and successfully predict binding at motifs for 38 TFs in a lymphoblastoid cell line, achieving superior classification performance compared to existing frameworks. Next, we implement Grad-CAM^25^, a post-analytical method for neural networks, and compute importance scores for the context regions of the binding motifs at single base pair resolution. These scores are further applied to uncover known and novel determinants of TF binding. Finally, we demonstrate that context bases predicted to be the most important binding determinants are under increased purifying selection compared to nearby bases. These important bases are enriched in disease-associated variants identified by genome-wide association studies. Overall, our framework enables novel insights into sequence features determining TF binding and identifies specific non-coding variants with potential disease relevance. To facilitate these and other applications, all variant level scores as well as our model framework, named AgentBind, are available at https://github.com/Pandaman-Ryan/AgentBind.

## Results

### Predicting binding status of transcription factor motif occurrences

We focused on 38 TFs active in the lymphoblastoid cell line GM12878 with ChIP-seq datasets available from the ENCODE Project^26^, and binding site motifs available from JASPAR^27^ (**Supplementary Table 1**). For each TF, we scanned the human reference genome (hg19) to identify all instances of its binding motif, which we refer to as the *core motif*. We extracted 1 kb genomic sequences centered on each core motif instance and labeled each sequence as bound (positive set) vs. unbound (negative set) based on the overlap of the core motif region with binding sites identified by ChIP-seq (**Methods, Fig. 1a**). For each TF, we chose an approximately equal number of positive and negative samples for analysis. On average for each TF, we obtained 18,892 sequences as input to our classification and interpretation modules.

**Figure 1:**
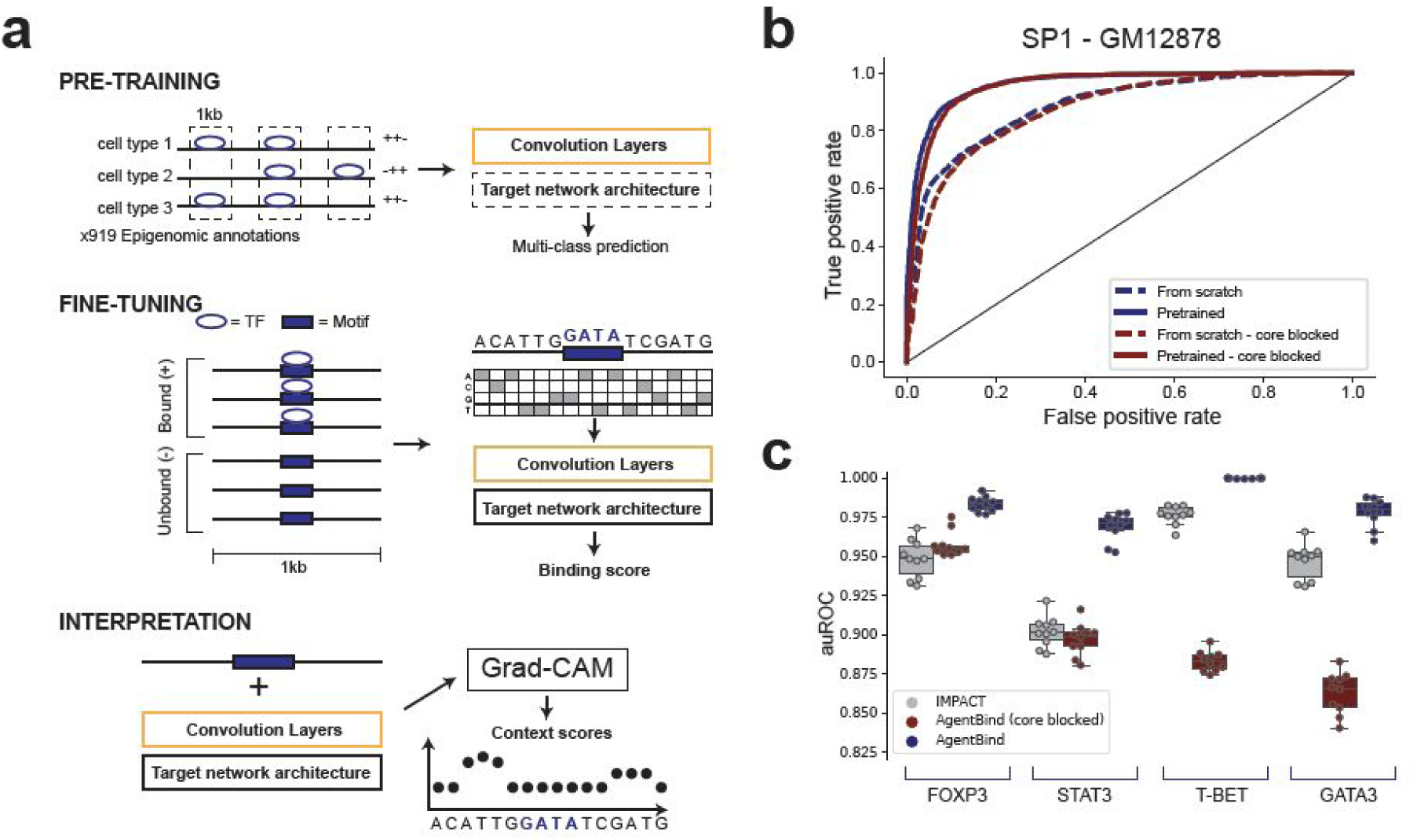
Overview of the AgentBind framework. **(a) Schematic of AgentBind**. In the pre-training step (top), AgentBind trains a convolutional neural network on ChIP-seq, DNAseI-seq, and other epigenomic annotations profiled in a variety of cell types. In the fine-tuning step (middle), AgentBind trains on sequences containing a core motif (purple box) for a TF of interest that are either bound (+) or unbound (-) to specifically learn context features determining binding status. In the interpretation step (bottom), Grad-CAM is used to score the contribution of each nucleotide to the binding prediction. **(b) Pre-training improves prediction of motif binding status**. A receiver operator curve (ROC) is shown for the TF SP1 in GM12878. Dashed lines show performance without the pre-training step (from scratch). Solid lines show performance using the pre-trained model. Dark red and blue lines show performance for models with and without the core motif blocked, respectively. **(c) Comparison to IMPACT**. For four TFs (FOXP3, STAT3, T-bet, and GATA3), we trained AgentBind to distinguish bound vs. unbound motifs and compared performance to IMPACT, which uses epigenomic features but not sequence information. Boxplots show the distribution of auROC values for 10 rounds of randomly selecting training (80%) vs. testing (20%) motif instances. Gray=IMPACT, dark blue=AgentBind without the core motif blocked, dark red=AgentBind with the core motif blocked.

The sizes of input datasets varied widely, from 182 to 107,539 total sequences per TF (**Supplementary Table 2**). To enable our framework to accommodate smaller datasets, we applied a two-step transfer learning scheme, including a pre-training step and a fine-tuning step. Transfer learning has been shown to dramatically reduce the amount of training needed for related classification tasks and improves the overall predictive performance compared to training from scratch^28^. In the pre-training step, we trained a CNN on 4,863,024 1 kb sequences annotated with a total of 919 ChIP-seq and DNase-seq profiles collected from ENCODE^26^ and the Epigenomics Roadmap Project^29^ across dozens of cell types (**Methods**). This step allows the CNN model to capture the common DNA patterns in regulatory regions and encode them in its convolutional layers. In the fine-tuning step, we customized an individual classification model for each TF to identify bound vs. unbound sequences as described above. Instead of training from scratch, in this step we initialized the convolutional layers with the parameters learned in the pre-training, such that our model could make use of the sequences features learned previously and focus on the novel context patterns specific to the TF of interest.

Our framework, AgentBind, is compatible with theoretically any CNN-based architecture. As examples we evaluated its performance using two popular neural network architectures, DeepSEA^18^ and DanQ^20^, separately. The DeepSEA architecture consists of three CNN layers and one fully connected layer. DanQ is a hybrid neural network architecture containing both CNN and recurrent neural network (RNN) layers. For each evaluation experiment, we reserved sequences from chromosome 8 for cross-validation and from chromosome 9 for testing in both the pre-training and fine-tuning steps. We applied two metrics to quantify the framework performance including area under the receiver operating characteristic curve (auROC) and the precision recall curve (auPRC). Average performance across all TFs was high (auROC 0.945 and 0.932 for DanQ and DeepSEA architectures, respectively), suggesting that binding is primarily controlled by local sequence features within a few hundred base pairs of the core motif. Notably, model performance was correlated with the difference in GC content between bound vs. unbound regions (Pearson r=0.58; p=0.00012; **Supplementary Fig. 1**), suggesting that for a subset of TFs GC content is a major predictor of binding status, in concordance with previous literature^12^.

In both evaluation experiments, transfer learning noticeably improved the predictive performance compared with models trained from scratch (**Fig. 1b**), especially for TFs with low sample sizes. For example, using our framework with DanQ, we achieved an auROC of 0.920 and auPRC of 0.882 for FOS, which contains only 182 sequences, whereas the same model trained from scratch failed to classify bound vs. unbound sequences (auROC=0.580, auPRC=0.543). Our framework, which focuses specifically on context regions by conditioning on presence of the core motif in both positive and negative sequences for one TF at a time, consistently outperformed multi-class models for this classification task (38/38 for DanQ and 32/38 for DeepSEA architectures showed higher auROCs). Full evaluation results for each TF are available in **Supplementary Tables 3 and 4**.

To further determine whether our model is primarily learning sequence features from context regions rather than from core motifs, we repeated our analyses with the central core motif instances masked. In most cases prediction performance was unchanged or only slightly reduced (average auROC decrease 0.015 (DanQ) and 0.029 (DeepSEA); **Supplementary Tables 5; Methods**). Overall this demonstrates our predictions are based primarily on context sequences. CTCF was a notable exception (auROC=0.94 and 0.80 before and after blocking, respectively), suggesting its bound vs. unbound regions have key differences within core motifs despite being similarly scored by position weight matrices (PWMs).

AgentBind automatically captures features determining binding status based on DNA sequence alone. We compared our results with the recently developed IMPACT method^30^, which tackles a similar classification task but instead using a broad range of manually extracted epigenomic features including histone modifications, TF binding, ATAC-seq, and DNAse-Seq profiles. We benchmarked each method on 4 TFs active in CD4+ T cells and applied the same training scheme as was used in the IMPACT study. AgentBind demonstrated higher auROC than IMPACT in all four cases (**Fig. 1c, Supplementary Table 6**). This suggests that the majority of determinants of binding for these TFs can be extracted directly from local sequence features. For FOXP3 and STAT3, performance was comparable to IMPACT even with core motifs blocked and making classification decisions purely based on surrounding sequence contexts. For T-BET and GATA3, AgentBind’s performance gain was partly driven by differences in the core motifs themselves.

### Identifying the context-specific determinants of TF binding

Although deep neural networks achieve high classification accuracy, compared to simpler linear models they are not trivially interpretable. Several techniques, including *in silico* mutagenesis^18,21,22^, DeepLIFT^25,31^, and saliency maps^32^, have previously been applied to interpret CNN results on DNA sequences. These methods quantitatively annotate the contribution of each nucleotide in a sequence toward the classification prediction. However, these methods face important disadvantages: *in silico* saturated mutagenesis is computationally inefficient as each perturbation requires a separate forward propagation through the network; DeepLIFT doesn’t support hybridized architectures containing RNNs, such as DanQ. Saliency maps are a group of promising methods that compute gradients of neural network outputs with respect to each nucleotide. Saliency map methods (1) require only one step of forward propagation per sample, (2) can be applied to any type of neural network, and (3) can be implemented easily under deep learning frameworks such as Tensorflow and PyTorch^33,34^. However, naive implementations (e.g.^32,35^) are highly sensitive to noise and are susceptible to model saturation^25,31^. Here, we applied and implemented Grad-CAM^25^, an advanced version of saliency maps, which overcomes this challenge using an aggregated distribution map of important sequence features and integrates this distribution map with input layer gradients through element-wise multiplication **(Methods)**. This method is computationally efficient and has been proven to be more stable than the vanilla saliency maps in many computer vision applications^25^.

We first designed a simulation experiment to evaluate the applicability of these model interpretation methods to identifying important context sequence features. We adapted a previously published evaluation scheme^36^, and generated a binary dataset with TF motifs embedded at known positions (**Methods**). Both positive samples and negative samples are 1kb long with GATA1 motifs embedded in the center. In each positive sample, we further randomly embedded 1-3 instances of the TAL1 motif in the context regions. This simulated binary dataset was fed into our framework and individual nucleotides for each sequence were annotated by each interpretation method. For this and downstream visualization analyses, we chose to focus on the DanQ architecture due to its superior performance in predicting binding affinity (**Supplementary Tables 3-4**). We computed signal-to-noise ratios for each method by taking the ratio of context scores in the embedded TAL1 motif regions to context scores in background regions. Grad-CAM greatly outperformed alternative methods (mean signal to noise ratio 99.0, 33.5, and 4.6 for GradCAM, vanilla saliency map, and saturated mutagenesis respectively; **Fig. 2a**) and could more precisely identify the embedded context motifs (**Fig. 2b**).

**Figure 2:**
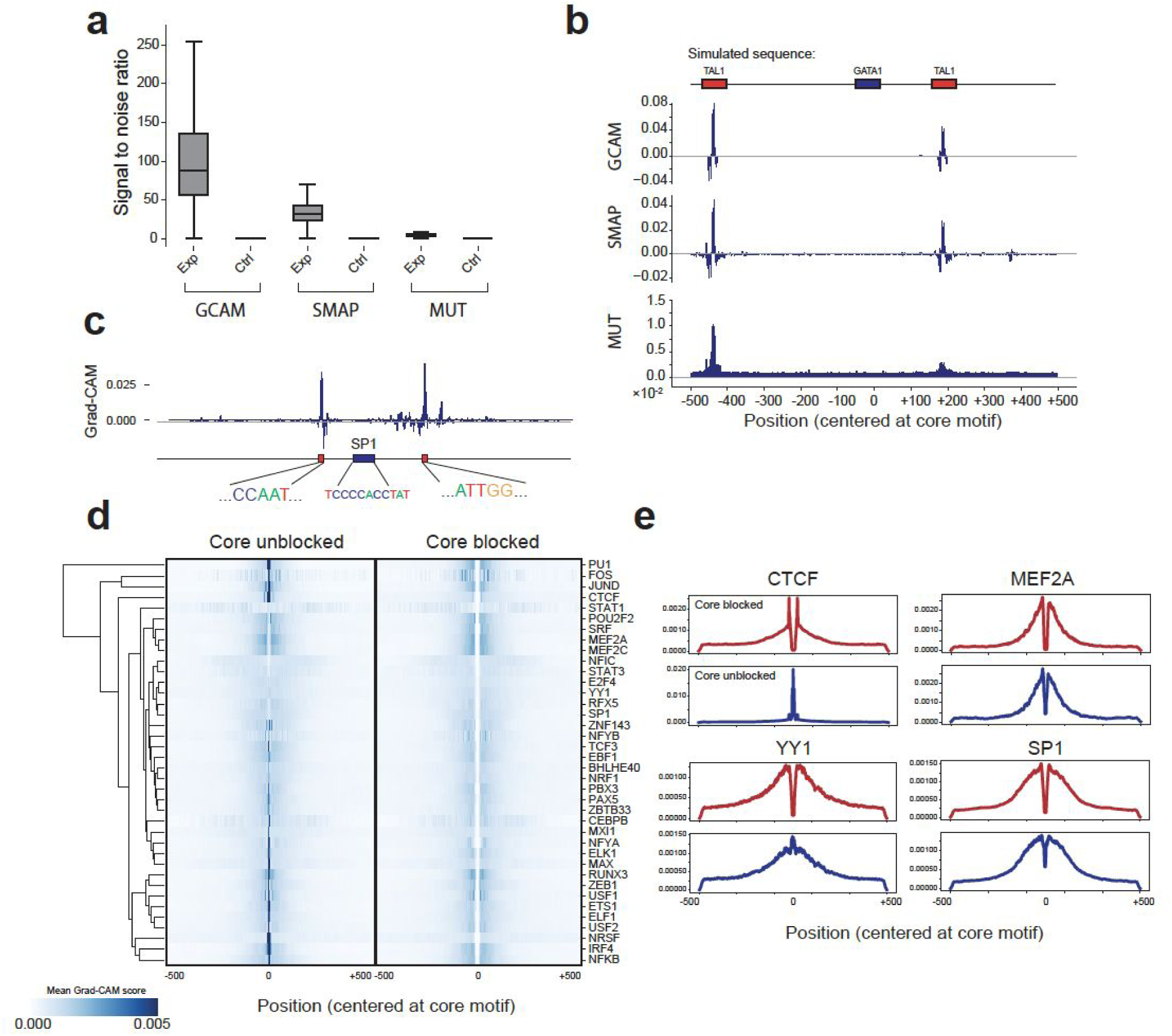
Interpreting the context-specific determinants of TF binding. **(a) The distribution of signal to noise ratio for three model interpretation methods**. The signal to noise ratio is computed as the mean score of bases within simulated TAL1 motifs compared to the mean score of background bases in the region. Control distributions were obtained by randomly sampling 1000 regions with the same size as the TAL1 motif. **(b) Example importance scores for a simulated region**. The top shows an example simulated sequence, with a central GATA motif and two context TAL1 motifs. Importance scores are shown for Grad-CAM (top-GCAM), saliency maps (middle-SMAP), and saturated mutagenesis (bottom-MUT). Note, by visualizing the annotation scores, we found that in some cases tall peaks were flanked by negative scoring regions. These negative regions are likely artifacts and do not correspond to regions negatively contributing to binding prediction. **(c) Example Grad-CAM scores for a region containing an SP1 motif**. The y-axis shows the Grad-CAM score of each nucleotide. Sequences are shown for the central SP1 motif and two regions with high Grad-CAM scores corresponding to NFY motifs. **(d) Aggregate Grad-CAM score profiles for each TF**. For each TF, we computed the average Grad-CAM score per position in positive sequences using either models with the core motif unblocked (left) or blocked (right). Values shown are Z-normalized across rows. **(e) Example aggregate Grad-CAM profiles**. For four representative TFs, average Grad-CAM scores are shown for models with the core motif blocked (top; dark red) or unblocked (bottom; dark blue).

We used GradCAM, the highest performing method, to compute context scores for each position in all bound (positive) input sequences for each of the 38 TFs described above. Score profiles for individual sequences highlight specific regions contributing to the classification. For instance, **Fig. 2c** shows an example interpretation of a weak SP1 binding at chr1:12289432-12290432. Grad-CAM scores identify two key regions approximately 100bp upstream and downstream of the core motif, respectively. These regions consist of the NFY motif (“CCAAT/ATTGG”), with which SP1 is known to co-bind^10^. Aggregating scores across all input sequences for each TF shows as expected that sequences closest to the core motif tend to have the highest impact (**Fig. 2d**). This pattern is present in models with and without the core motif masked. However, aggregate score profiles differ noticeably for different TFs. For example, whereas the important bases for predicting CTCF binding are highly concentrated directly adjacent to the motif, important bases for YY1 are spread across the entire 1 kb region (**Fig. 2e**). In concordance with our results above, differences in core motifs themselves receive high importance scores for some TFs (e.g. PU1, CTCF) but not others (e.g. MEF2A, SP1, **Fig. 2e**).

### Grad-CAM scores give insight into features of TF binding

We next sought to use Grad-CAM score profiles to identify context sequence features with strongest impact on binding status for each motif. To this end, we segmented the positive (bound) input sequences into subsequences of length 5 using a sliding window and identified 5-mers with highest average Grad-CAM scores, which we denote as top 5-mers (**Methods**). For each input sequence, we only scored the strand (forward or reverse complement) matching the core motif. We then performed a Fisher’s Exact test to determine whether each possible 5-mer sequence was enriched among top 5-mers. We performed hierarchical clustering on the matrix of odds ratios for each 5-mer sequence+TF combination to identify key context features determinant of binding for one or more TFs (**Fig. 3a**). Notably, multiple related 5-mers cluster tightly and can be readily assembled into longer enriched motifs.

**Figure 3:**
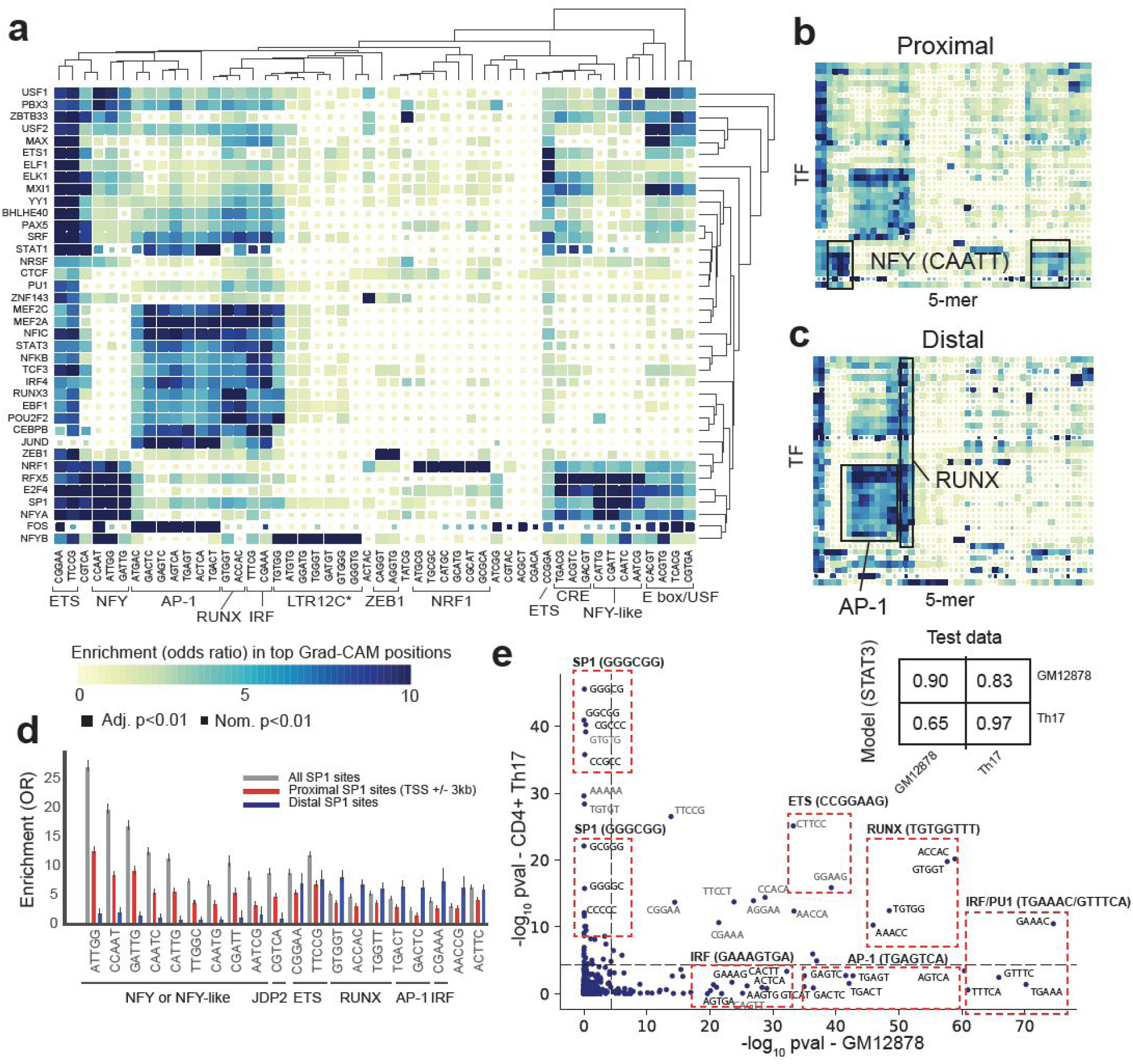
Identifying key context sequence features for TF binding in GM12878. **(a) Enrichment of 5-mers in most influential context regions**. The heatmap shows the enrichment of each sequence in regions with the highest Grad-CAM scores for each TF. (**Methods**) Colors denote odds ratios and the size of boxes denote statistical significance. Dendrograms were computed using hierarchical clustering on the odds ratio matrix. Boxed and annotated 5-mers correspond to known core motifs. **(b) Enrichment of 5-mers in high-scoring Grad-CAM regions for proximal binding sites**. Proximal TF binding sites are defined as being within 3kb of a transcription start site. Rows and columns are ordered the same as in **(a). (c) Enrichment of 5-mers in high-scoring Grad-CAM regions for distal binding sites**. Distal binding sites are defined as being at least 3kb from the nearest transcription start site. Rows and columns are ordered the same as in **(a). (d) Comparison of top scoring 5-mers in proximal vs. distal SP1 sites**. Bars show the odds ratio of enrichment of each sequence in top 5-mers for all (gray), proximal (red) and distal (blue) SP1 sites. The 5-mers with the top 20 odds ratios across all sites are shown. Error bars show 95% confidence intervals on odds ratios. **(e) Cell-type specific enrichment of 5-mers influential for STAT3 binding**. Enrichments were computed as in **(a)** using models trained separately in either GM12878 or CD4+ Th17 cells. The inset shows the auROC for each model evaluated on each of the two cell types. The main plots shows the −log10 p-value of enrichment of each 5-mer in GM12878 (x-axis) vs. CD4+ Th17 cells (y-axis). Each dot represents a single 5-mer. Dashed lines denote the cutoff for statistical significance (p<0.05) after correcting for the number of kmers tested.

The 5-mers predicted to have the highest impact on TF binding based on Grad-CAM scores recapitulate multiple known determinants of binding for these TFs. First, the top scoring sequences for a TF often closely match the core motif of the TF itself. For example, the presence of an additional NFY motif (5’-CCAAT-3’) in the context is a strong determinant of binding of NFYA (p<10e-200; OR=23.7) and NFYB (p<10e-200; OR=10.9). Similarly, kmers from the NRF1 (5’-TGCGCATGCGCA-3’) and ZEB1 (5’-CAGGTG-3’) motifs score highly for NRF1 (p<10e-200; OR=14.1 for ATGCG) and ZEB1 (p<10e-200; OR=18.1 for CAGGT), respectively. This result is consistent with previous literature suggesting that increasing the number of localized motifs for a particular TF (referred to as homotypic clusters) can promote binding^37,38^, a trend that has already been reported for these particular TFs^37,38^.

Second, we identify distinct clusters of TFs for which binding of the core motif is predicted by similar motifs shown to interact previously with those factors. For example, the NFY motif (5’-CCAAT-3’) is a strong determinant of binding for factors NFYA, NFYB, SP1^39^, E2F4^40^, and RFX5^41^. In another cluster, the AP-1 motif (5’-TGA G/C TCA-3’), bound by a dimerization of JUN and FOS^42^, is predicted as a strong determinant of binding for a variety of TFs, including FOS and JUN, as well as other known co-binders, including CEBPB^43^, MEF2A^44^, and IRF4^45^. The RUNX motif (5’-TGTGGT-3’) is a similarly strong determinant of binding in this cluster. In the third cluster, the ETS motif (5’-CGGAA-3’) is a determinant of binding for MXI1, ELF1, ELK1, ETS1, and other factors. Both AP-1 and NFY have previously been shown to act as pioneer factors^46,47^, consistent with the observation that their motifs are predictive of binding for a wide range of factors. Furthermore, both NFY and ETS motifs have been shown to act as cardinal elements of promoter regions^48^, which dictate specific signatures of co-factors at largely distinct sets of promoters.

Grad-CAM scores captured several unexpected enrichments. For example, subsequences comprising the sequence 5’-GGGTGGGTATGTG-3 were strongly enriched in top 5-mers for NFYB, which forms a trimeric complex with NFYA and NFYC and should primarily recognize the NFY motif 5’-CCAAT-3’. Further inspection of specific NFYB-bound regions containing this sequence showed all consist of the long terminal repeat (LTR) element LTR12C, which has previously been reported to contain NFYB binding sites^49^. Additionally, we found that while binding of NFYA, NFYB, and factors with similar binding patterns is strongly predicted by the presence of canonical NFY motifs in the context, 5-mers similar to CCAAT/ATTGG but mismatched at one nucleotide also received high Grad-CAM scores (**Fig. 3a, Supplementary Fig. 2**). In some but not all cases, these weakly match extended motifs for NFYA or NFYB (e.g. CCATT, CCGAT). To determine whether this observation could be an artifact due to our model computing similar scores for related kmers, we examined enrichment of other sets of 5-mers different by a single nucleotide from a canonical motif, which did not recapitulate the trend observed for NFY-like kmers (**Supplementary Fig. 2**). These results are consistent with a model by which NFY-like sites nearby true CCAAT boxes promote binding, perhaps by helping guide TFs toward the correct binding site.

Finally, we hypothesized that the sequence context features promoting binding of a particular TF to its core motif might differ between TFs in promoter regions (+/-3kb from transcription start sites [TSS], denoted as “proximal”) vs. “distal” regions (>3kb from the nearest TSS). We repeated our analysis of top 5-mers separately for proximal and distal binding sites (**Fig. 3b-c**). Overall, NFY and NFY-like motifs, which are known cardinal elements of promoter regions^48^, have a much stronger influence on proximal binding sites, but very little influence on distal sites. We further examined these enrichments specifically for SP1 (**Fig. 3d**), for which the most influential 5-mers differed dramatically between proximal and distal sites. Similar to the trend observed overall, NFY sites have strong influence on proximal sites and almost none on distal sites. ETS motifs have strong and similar impacts on both proximal and distal sites. On the other hand, RUNX, AP-1, and interferon regulatory factor (IRF) motifs have strong influence only on distal sites. Overall, these results suggest the context features influencing binding for SP1 and other factors are largely orthogonal at promoter vs. enhancer regions and are likely governed by separate sets of pioneer factors.

### Cell-type specific models detect distinct TF binding determinants

TF-binding is well known to be highly cell-type specific. Thus we hypothesized that CNN models trained for the same TF but on different cell types would capture cell-type specific regulatory features. We trained a separate model based on the DanQ architecture to classify STAT3 binding sites as bound vs. unbound in GM12878 and CD4+ Th17 cells. We then used each model to predict STAT3 binding in each of the two cell types. As expected, models trained and evaluated on the same cell type showed superior performance (auROC=0.9 and auROC=0.97 for the GM12878 and CD4+ models, respectively) compared to models evaluated on orthogonal cell types (auROC=0.65 using the CD4+ model to predict GM12878, and auROC=0.83 using the GM12878 model to predict CD4+) (**Fig. 3e**).

We then computed Grad-CAM scores for each sequence predicted to be bound and identified sequences enriched in the top 5-mers from each model as described above. Several 5-mers, such as those belonging to ETS and RUNX motifs, are significant (adjusted p<0.05) for both cell types. However, the majority of top scoring 5-mers are distinct for each cell type (**Fig. 3e**). For GM12878, IRF, and AP-1 motifs are most strongly predictive of STAT3 binding. Both IRFs and AP-1 have been previously shown to be stimulated by the LMP1 protein active in Epstein-Barr virus used to immortalize LCLs^50,51^. On the other hand, CG-rich sequences, including SP1 binding motifs, are most influential for STAT3 binding in CD4+ Th17 cells. SP1 is known to recruit the SWI/SNF chromatin remodeling complex and thus may help in making DNA more accessible for binding of other factors. Overall these results are consistent with a model whereby STAT3 binds to already accessible regions^52^, which were made accessible by different factors in each cell type. Our model readily captures these cell-type specific context features.

### Grad-CAM scores identify variants under negative selection

A major result arising from genome-wide association studies (GWAS) has been that the majority of genetic variants impacting complex human traits fall in non-coding regions^53^. It is thought that most of these variants act primarily by affecting gene regulation through TF binding or other mechanisms^54^. It has further been demonstrated that core TF binding sites are under weak but significant purifying selection^55^, consistent with their weak but prevalent effects on medically relevant traits. We reasoned that context positions identified as most influential for TF binding would be similarly under increased purifying selection compared to surrounding genomic regions.

To quantify selection for a set of genomic positions, we assessed whether those positions are depleted of common genetic variation compared to nearby positions. We focused on single nucleotide polymorphisms (SNPs) present in the gnomAD v2 database^56^ overlapping sites that were scored by Grad-CAM for each TF, and computed the percentage of SNPs for which the alternate allele is observed only once (termed singletons) indicating the variant is extremely rare. This “percent singleton” metric has previously been used as a proxy for deleteriousness of a set of SNPs^57^. When aggregating across all TFs, the top-scoring 1% of positions based on Grad-CAM scores show a significantly higher singleton percentage compared to all sites scored by Grad-CAM (p=2e-13). When analyzing TFs individually, top-scoring positions have increased singleton percentages for 31/38 TFs (**Fig. 4a**), significantly greater than the 50% expected by chance (p=6e-5). For multiple TFs (16 out of 38), influential context regions showed higher singleton rates than the core TF motifs themselves (**Supplementary Fig. 3**).

**Figure 4:**
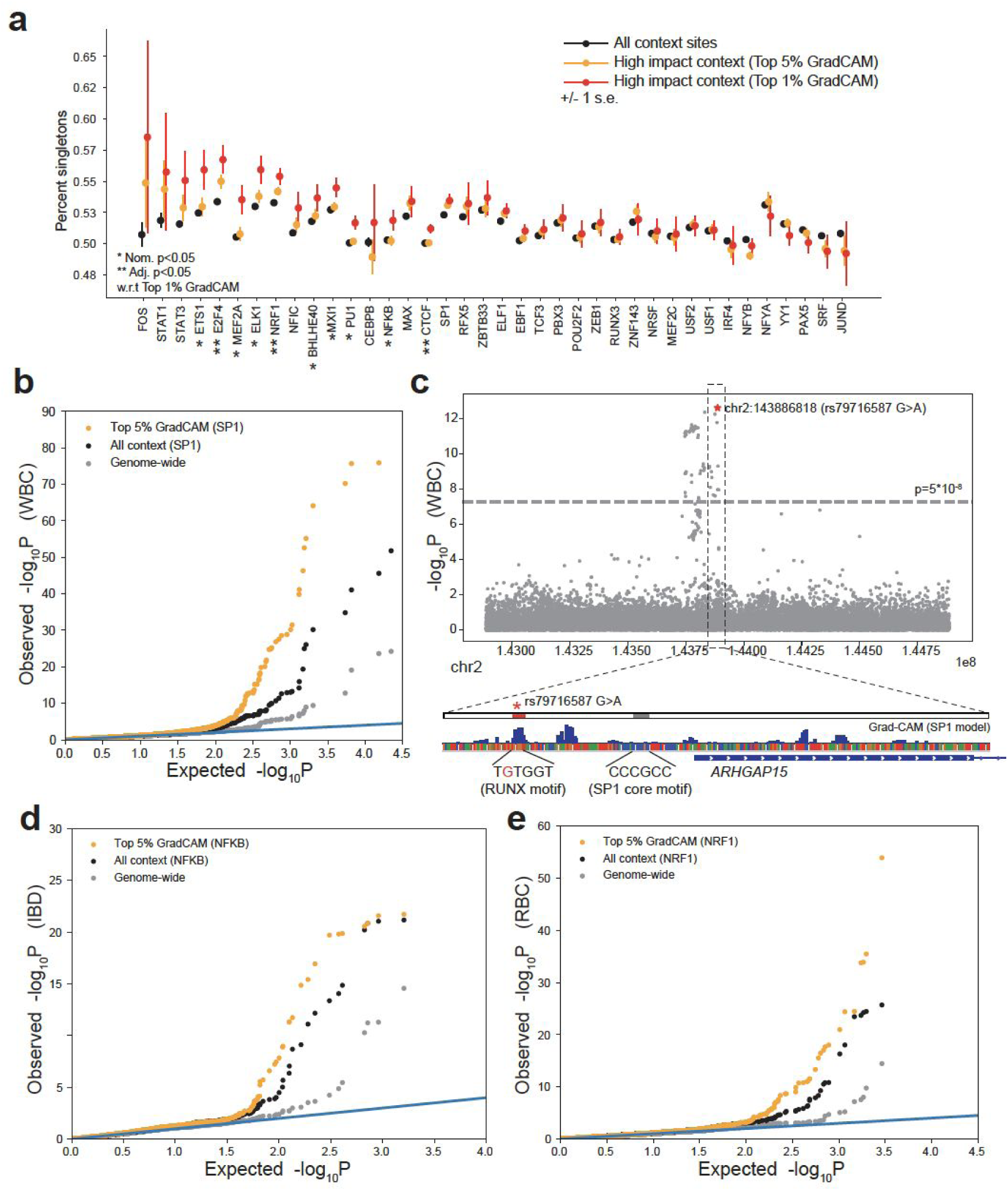
Grad-CAM scores prioritize high-impact context SNPs. **(a) Singleton rate of context SNPs**. The plot shows the percent of SNPs in each category that are singletons. Black=all context sites, orange=context sites with top 5% Grad-CAM scores, red=context sites with top 1% Grad-CAM scores. Error bars show +/- 1 s.e. **(b) Quantile-quantile plot of association with white blood cell count (WBC)**. For each category of SNPs, the plot shows the distribution of p-values for association with WBC. Gray=randomly selected genome-wide SNPs, black=randomly selected context SNPs, orange=top 5% Grad-CAM sites for SP1. Randomly selected SNPs were chosen to match the Grad-CAM sites on allele frequencies. The blue diagonal line gives the expected uniform distribution. **(c) Example association of a top-scoring Grad-CAM SNP for SP1 with WBC**. In the top plot, the x-axis gives the genomic location and the y-axis gives the −log10 p-value of association with WBC for each SNP. The gray horizontal line gives the genome-wide significance threshold (5×10^−8^). The SNP denoted with the red star is the top ranked Grad-CAM SNP for SP1 binding in this region. The bottom plot zooms in on this region. The candidate SNP (rs7916587:G->A) disrupts a RUNX binding site in the context of an SP1 binding site near the transcription start site of *ARHGAP15*. **(d) Quantile-quantile plot of association with inflammatory bowel disease (IBD)**. Same as **(b)** except p-values give associations with IBD and data is shown for the NFKB model. **(e) Quantile-quantile plot of association with red blood cell count (RBC)**. Same as **(b)** except p-values give associations with RBC and data is shown for the NRF1 model.

We tested whether high-scoring context bases are enriched for SNPs that are contributing to medically relevant human traits. To this end, we integrated our Grad-CAM results for each TF with publicly available GWAS SNP summary statistics for a variety of blood-related and other traits (**Methods**). We then examined the distribution of association p-values for SNPs with high Grad-CAM scores compared to other variant classes. In this experiment, the p-value distributions most strongly inflated compared to the expected uniform distribution indicate an enrichment for trait-associated SNPs.

As a test case, we examined SNPs in context regions (+/- 500bp from the core motif) of multiple TF binding sites in GM12878 and their associations with white blood cell count (WBC) measured by GWAS in the UK Biobank cohort^58^ (**Supplementary Fig. 4**). The results demonstrated that SNPs near SP1 binding sites have most strongly inflated p-values for WBC. Restricting to SNPs at positions with the top 5% Grad-CAM scores further inflated the p-value distribution (**Fig. 4b**), indicating the strongest enrichment for WBC association within the most influential context bases. We examined one example SNP from this distribution, rs79716587:A>G, with a high Grad-CAM score (greater than the scores of 99.5% positions) and strongly associated with WBC (p=2.4e-13). This SNP is the top-ranked GWAS SNP in the region and disrupts a RUNX motif in the context of an SP1 binding site (**Fig. 4c**). It falls within the core promoter of *ARHGAP15*, which has previously been identified as a master negative regulator of neutrophil function in mice^59^. We repeated this analysis for a variety of TF-trait pairs (**Supplementary Fig. 4**) including NFKB, a TF known to be involved in regulating immune response^60^, in inflammatory bowel disease^61^ (**Fig. 4d**) and NRF1, a TF with known involvement in heme synthesis^62^, in red blood cell count^58^ (**Fig. 4e**). We consistently observed that SNPs near binding sites of key TFs are enriched for associations, and that the highest ranked Grad-CAM sites have most strongly inflated p-values compared to other context SNPs. These results suggest that context bases most influential for TF binding, beyond core TF motifs themselves, play a critical role in a variety of human traits.

## Discussion

Here we present AgentBind, a machine learning framework, to predict whether particular instances of TF motifs in the genome are bound vs. unbound in a given cell type and to identify context features most important for determining binding. We tested our framework on simulated datasets and applied it to identify and interpret binding sites for 38 TFs profiled by ENCODE in the lymphoblastoid cell line GM12878. Remarkably, our framework shows that binding status at motifs for most TFs analyzed can be predicted almost entirely (mean auROC=0.94) by 1 kb of local sequence context alone. Applying Grad-CAM, a state of the art model interpretation technique, pinpoints the most influential context bases. We demonstrate that these scores can be used to identify key context features for each TF, find specific bases under purifying selection, and to enrich for likely causal GWAS variants in non-coding regions for various traits. Notably, while we focused on TFs in GM12878 using the DanQ architecture, this framework can similarly be applied to any TF and cell type of interest for which ChIP-sequencing data is available and a flexible range of CNN model architectures.

One major application of our framework is to learn which context sequences beyond core motifs themselves contribute to binding. By obtaining nucleotide-resolution annotation of important bases, we are able to identify specific kmers with highest influence. Notably, these kmers are not always the most frequent kmers in positive vs. negative examples, and thus can only be identified using nucleotide-level, rather than region-level, scores. Our analysis of most influential 5-mers for a group of well-studied TFs in GM12878 identified many known TF relationships, such as co-binding of NFY and SP1, the role of AP-1 as a pioneer factor, and the presence of an NFYB motif embedded in an LTR12C repeat element. We applied a similar analysis of STAT3 binding in GM12878 vs. CD4+ Th17 cells and found that the kmers identified can be strikingly different across different cell types. Our framework can now easily be applied to new sets of TFs and cell types to uncover novel grammars for less well-characterized TFs.

Another major insight is that important context bases are under stronger negative selection than surrounding context bases based on analysis of SNP allele frequencies in gnomAD. In some cases, selection was stronger at context bases than for core motifs themselves, which have previously been shown to be under purifying selection^55^. This result validates that Grad-CAM scores are picking up important non-coding positions that are more likely to have consequences for human health. Indeed, integration of our nucleotide-level scores with published GWAS summary statistics for blood-related and autoimmune traits shows that most influential context bases for relevant TFs are enriched for association compared to other context bases or genome-wide SNPs, indicating the Grad-CAM or similar scores will be helpful metrics to interpret and prioritize non-coding variants contributing to complex traits.

Our study faced several limitations: (***i***) modifications to the training process are likely to improve performance. For example, we chose to use 1 kb context regions centered on core motifs, as this size of genomic region has been shown to work well in predicting general TF binding. While 1kb of context provides remarkable predictive power, it is likely that some aspects of TF binding are controlled by more distal features. Further, our current training procedure attempts to classify binding status of all core motif instances of a particular TF. However, our results examining important predictive 5-mers in proximal vs. distal regions suggests binding in these regions might be controlled by separate mechanisms and thus may better be modeled separately. (***ii***) Model interpretation techniques can be further improved to extract more complex rules for TF binding. While our kmer analysis of top scoring Grad-CAM regions is helpful for interpreting specific context motifs contributing to binding of each factor, it does not readily reveal rules such as motif spacing, orientation, and combinations. These rules are likely learned by CNN models but are not captured by nucleotide-level scores. (***iii***) Finally, while our method accurately prioritizes specific high-impact variants under negative selection, the current implementation only computes scores for a small subset of all non-coding variants. In particular, it requires deciding on a TF and cell type of interest *a priori* and computes scores only for SNPs within 500bp of motifs for that TF. Thus it cannot serve as a standalone method for scoring all non-coding sites. In future work, we will evaluate combining scores across TFs to compute a composite score, and will evaluate the utility of these scores for variants not falling near binding sites.

Altogether, our study provides a valuable machine-learning framework for helping decode the rules by which TFs bind their target sites and identifying specific non-coding nucleotides with the strongest effects on binding. Example future applications of these results include: application to novel TFs and cell types to learn cell-type specific rules of additional TFs, studying signals of selection at TF binding regions, and integration of nucleotide-level scores into GWAS fine-mapping frameworks. To facilitate these and other studies, Grad-CAM scores for all TF models studied here and code for running these models on novel datasets are publicly available at https://github.com/Pandaman-Ryan/AgentBind.

## Online Methods

### ChIP-sequencing datasets and preprocessing

We used FIMO^63^ v4.12.0 to identify all instances of the motif for each TF across the human reference genome (hg19). Fimo takes the reference genome and target motifs for each TF as input and returns all occurrences of the target motif (using the default p-value threshold p<10^−4^). Motifs for each TF were obtained from Jaspar (**Supplementary Table 1**). We intersected motif instances with binding sites as identified by ChIP-sequencing available for each TF in GM12878 from the ENCODE Project^26^ (peak file accessions given in **Supplementary Table 1**) using a custom script. Motif instances (core motifs) were labeled as positive if they were fully within ChIP-seq peaks for the TF. All other instances were labeled as negative. We extended each core motif region to include 1 kb centered at the motif. For each sequence, we included it and its reverse complement sequence for the training procedure described below. In the experiments that required core motifs to be blocked, we substituted the motif region with a string of “N”‘s of the same length as the Jaspar motif.

The binary datasets we acquired above were highly imbalanced: the negative samples are 433 times more than positive samples on average (**Supplement Table 2**). In order to balance the dataset ratio while alleviating effects of differences within the core motif, we chose an identical number of negative and positive samples for each TF while requiring the distribution of motif match p-values to be similar. To obtain p-value matched sets, we binned −log10 p-values of motif matches into bins of size 0.1. We then selected the same number of positive and negative samples from each bin.

### CNN model architecture and training

We implemented DeepSEA and DanQ architectures using Tensorflow^34^ v1.9.0. The well trained models and their associated code are available in our Github repository (https://github.com/Pandaman-Ryan/AgentBind). DeepSEA consists of three convolutional layers and two fully connected layers, and DanQ consists of one convolutional layer, one bidirectional recursive neural network layer and two fully connected layers. We applied sigmoid cross entropy as the loss function for both models.

The dataset used in the pre-training step was downloaded from DeepSEA’s website (http://deepsea.princeton.edu/media/code/allTFs.pos.bed.tar.gz). It contains 4,863,024 1000 bp sequences annotated with a total of 919 ChIP-seq and DNase-seq profiles. We leave out sequences on chromosome 8 for cross validation and sequences on chromosome 9 for testing. We applied one-hot encoding to convert nucleotides in sequences into 4-element vectors as has been done in previous studies^18,21^. “N”s were converted into vectors with entries of 0.25 for each of the four nucleotides. During training, we initialized all model parameters with random Gaussian noise with mean 0 and standard deviation 1e-2, and trained this model on the DeepSEA compendium dataset until the loss function converged. In the fine-tuning step, we used the same architectures as in the pre-training step, and we built an independent model for each TF of interest using the labeled dataset described in the previous section. From the pre-trained model, we transferred its convolutional layers and RNN layer into the new models, but initialized the fully connected layers again with random Gaussian noise. These new models were further fine-tuned on our TF binary datasets until convergence.

### Benchmarking experiment against IMPACT

The IMPACT study focused on TFs active in T cells and created their own binary (bound vs. unbound) datasets for TFs including FOXP3 (Treg), GATA3 (Th2), STAT3 (Th17) and T-BET (Th1). The coordinates of motif instances for these four TFs were published on the the IMPACT Github repository (https://github.com/immunogenomics/IMPACT). In our benchmarking experiment, we used an identical set of motif instances, extending them into 1 kb sequences to train our model.

We applied an identical training scheme as was used by IMPACT: we randomly selected 80% of the samples in the input dataset for training and tested on the remaining samples. We evaluated our method in four situations using different architectures and core motif treatments (DeepSEA or DanQ architecture, with core motif blocked or unblocked), and for each situation we conducted 10 parallel trials with different selections of the test set.

### Model interpretation simulation experiments

We simulated a binary dataset for evaluating interpretation methods consisting of 50,000 samples for training, 1,000 for cross validation and 1,000 for testing. Labels were assigned evenly with same number of positives and negatives. All the samples were sequences of length 1000 bp containing the GATA1 motif (http://compbio.mit.edu/encode-motifs/) in the center. Context bases were generated by sampling the nucleotides A, C, G, and T at each position with probabilities 0.3, 0.2, 0.2 and 0.3 respectively. The number of TAL1 motifs (http://compbio.mit.edu/encode-motifs/) embedded in positive samples followed a Poisson distribution but truncated after 3. No TAL1 motifs were embedded in negative samples. These simulated sequences were fed into our AgentBind framework with DanQ architecture and annotated at nucleotide resolution using the model interpretation methods described below.

### Model interpretation methods

We implemented three separate model interpretation techniques using the simulated data described above. Each of these methods computes individual scores for each nucleotide of the input sequence indicating its importance in determining the model’s prediction.

For in silico mutagenesis, we performed computational mutations to assess the importance of every base of the input sequences. More specifically, we substituted each base with its three possible nucleotide substitutions and recorded the changes made by them in terms of the output prediction scores. The greatest score change was used to represent the importance of this base.

For vanilla saliency maps, the importance of each base was quantified using the gradient of the output prediction score with respect to this base. This step was accomplished using a Tensorflow build-in function “gradients”.

In our implementation of Grad-CAM, we chose the first convolutional layer as the layer of interest. This layer contains distribution maps for various sequence features. Following the weighting method proposed by the Grad-CAM authors^25^, we quantified the importance of these sequence features and computed a weighted summation of all the distribution maps. In comparison with vanilla saliency map which evaluates the importance of each base individually, this aggregated map highlights the regions that are important to the binding activities. To combine the best aspects of these two maps, we then merged the aggregated distribution map with the vanilla saliency map through element-wise multiplication.

We annotated the samples labeled as positive using these interpretation methods and used these annotation scores for downstream kmer enrichment and singleton analyses.

### Kmer enrichment analysis

In this analysis, we first segmented all the input sequences into 5-bp long subsequences using a sliding window and removed subsequences overlapping with core motifs in the center. Next, for each subsequence, we quantified its importance by averaging the Grad-CAM scores of each base. For each factor, we ranked all the subsequences based on their Grad-CAM scores and marked the top 1% as top 5-mers.

We used a Fisher’s Exact test to determine whether each 5-mer was enriched among high-importance 5-mers for each TF. Fisher’s Exact tests were performed using the fisher_exact function in the stats module of the Python scipy library^64^ v1.3.1. 5-mers were matched to published motifs based on manual inspection.

### Singleton analysis

For each TF, we overlapped sequences bound sequences scored by Grad-CAM above with single nucleotide polymorphisms (SNPs) in the control samples reported in the gnomAD v2 dataset^56^. For positions overlapping gnomAD SNPs, we recorded observed counts of minor alleles. We then labeled sites where the minor allele counts were 1 as singletons. We only included samples annotated in gnomAD as healthy controls (n=5,442 individuals) in our analysis and required a minimum total allele count of 1000. Sites not overlapping a gnomAD SNP (i.e. minor allele count of 0) were excluded from singleton analysis. The singleton ratio of a group of sites is then simply defined as the percentage of SNPs in that category that are singletons.

We computed singleton rates for: all context SNPs, core motif regions, and top-scoring positions. Here, top-scoring positions are defined as positions with highest Grad-CAM scores within each sequence. In this experiment, we calculated singleton rates respectively for the positions with top 1% Grad-CAM scores and top 5% Grad-CAM scores in each input sequence.

### Integration of Grad-CAM and GWAS summary statistics

Blood trait summary statistics^58^ were obtained from ftp://ftp.sanger.ac.uk/pub/project/humgen/summary_statistics/human/2017-12-12/. We used the files wbc_N172435_narrow_form.tsv.gz and rbc_N172952_narrow_form.tsv.gz for WBC and RBC respectively.

IBD summary statistics were downloaded from the International Inflammatory Bowel Disease Genetics Consortium (IIBDGC) website. We used the file EUR.IBD.gwas_info03_filtered.assoc with summary statistics in Europeans (ftp.sanger.ac.uk/pub/consortia/ibdgenetics/iibdgc-trans-ancestry-filtered-summary-stats.tgz).

As in the singleton analysis, we defined the top-scoring positions are defined as positions with highest Grad-CAM scores within each sequence.

## Acknowledgements

This study was supported in part by NIH/NHGRI 1R21HG010070-01 (M.G.), the Microsoft Genomics for Research program, and an Amazon Web Services research award. We thank NVIDIA for donating a Tesla K40 GPU to support this project. We additionally thank Chris Benner and Alon Goren for helpful comments.

## Author contributions

A.Z. designed and performed analyses and helped write the manuscript. M.L. and C.W. helped perform analyses. H.S. helped design the study. M.G. conceived the study, supervised analyses, and helped write the manuscript.

