## Supplementary Figures for "Deep neural networks identify context-specific determinants of transcription factor binding affinity"

**Supplementary Figure 1: Model performance related to GC content differences**

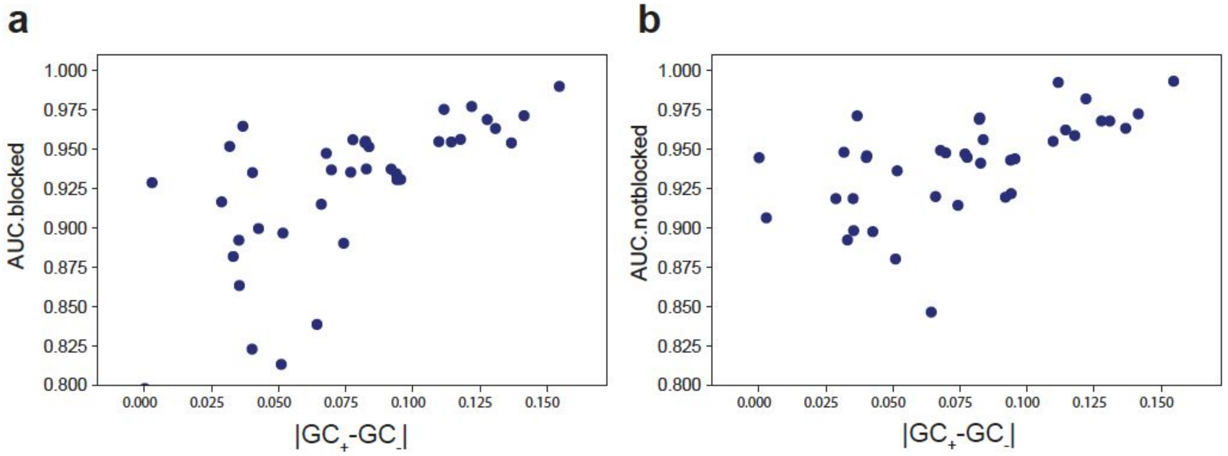


The x-axis shows the absolute value of the difference in mean GC content for positive (bound) vs. negative (unbound) sequences with the motif for each TF. The y-axis shows model performance. Each dot represents a single TF. Left=models with the core motif blocked; right=models with the core motif unblocked.

**Supplementary Figure 2: Enrichment of similar 5-mers in top scoring GradCAM positions**

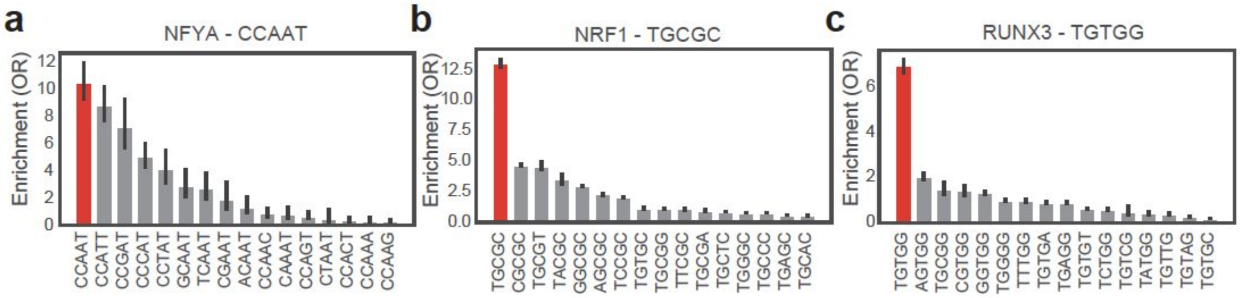


Red Bars show the odds ratio of enrichment of 5-mers matching canonical motifs for each factor in the top 1% of Grad-CAM sites. **(a)** “CCAAT” for NFYA, **(b)** “TGCGC” for NRF1, and **(c)** “TGTGG” for RUNX3. Gray bars show enrichments for all 5-mers that match the canonical motif at all except one position. Error bars show the 95% confidence intervals on odds ratios.

**Supplementary Figure 3: Singleton rate of context SNPs vs. core motif regions**

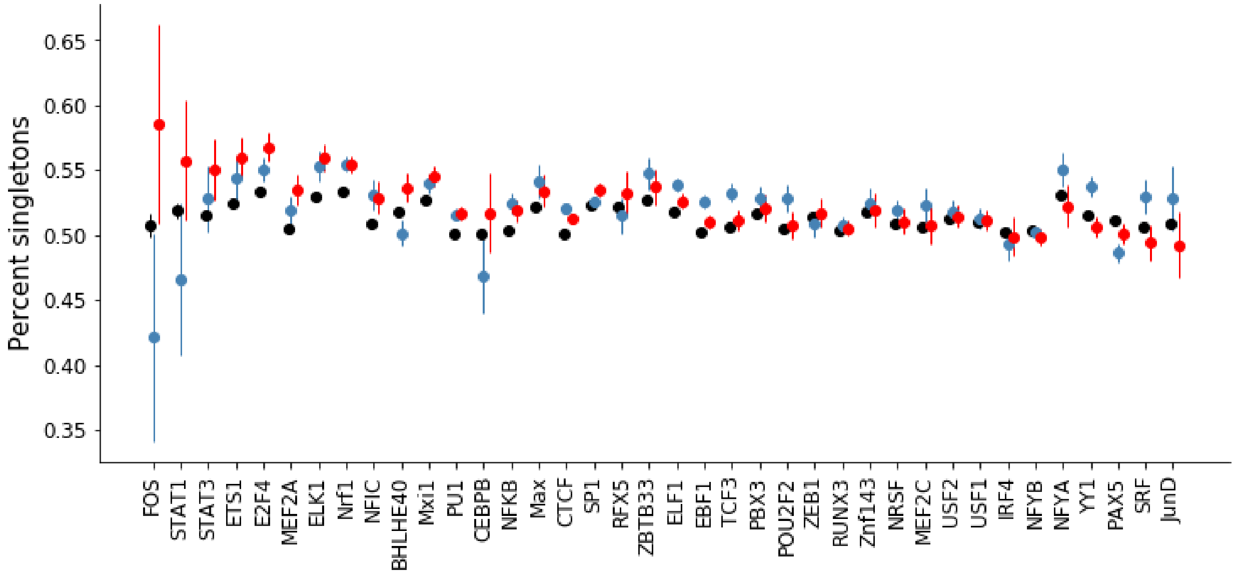


The plot shows the percent of SNPs in each category that are singletons. Black=all context sites, blue=core motif positions, red=context sites with top 1% Grad-CAM scores. Error bars show +/- 1 s.e.

**Supplementary Figure 4: High scoring context regions enriched for trait-associated SNPs**

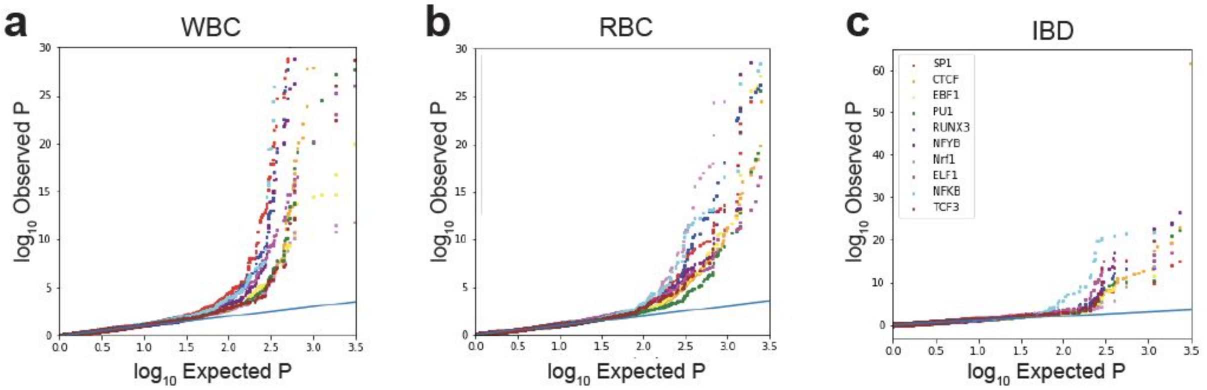


Quantile-quantile plots show the distribution of p-values for association with each trait (left=white blood cell count, middle=red blood cell count, right=inflammatory bowel syndrome,) for all SNPs within 1kb context regions of binding sites for each TF. Colors denote separate TFs as given in the legend in the figure.
